# VCF/Plotein: A web application to facilitate the clinical interpretation of genetic and genomic variants from exome sequencing projects

**DOI:** 10.1101/466490

**Authors:** Raul Ossio, Diego Said Anaya-Mancilla, O. Isaac Garcia-Salinas, Jair S. Garcia-Sotelo, Luis A. Aguilar, David J. Adams, Carla Daniela Robles-Espinoza

## Abstract

**Purpose:** To create a user-friendly web application that allows researchers, medical professionals and patients to easily and securely view, filter and interact with human exome sequencing data in the Variant Call Format (VCF).

**Methods:** We have created VCF/Plotein, a web application written entirely in JavaScript using the Vue.js framework, available at http://vcfplotein.liigh.unam.mx. After a VCF is loaded, gene and variant information is extracted from Ensembl, and cross-referencing with external databases is performed via the Elasticsearch search engine. Support for application-based gene and variant filtering has also been implemented. Interactive graphs are created using the D3.js library. All data processing is done locally in the user’s CPU to ensure the security of patient data.

**Results:** VCF/Plotein allows users to interactively view and filter VCF files without needing any bioinformatics knowledge. A number of features make it especially suited for the medical community, such as its speed, security, the ability to filter by disease or gene function, and the ease with which information may be shared with collaborators/co-workers.

**Conclusion:** VCF/Plotein is a novel web application that allows users to easily and interactively filter and display exome sequencing information, and that is especially suited for bench researchers, medical professionals and patients.

## INTRODUCTION

Exome sequencing (ES) has been highly successful at identifying genetic variation contributing to a large number of human phenotypes, from germline changes that underlie rare Mendelian disorders and complex diseases, to somatic mutations that drive carcinogenesis.^1,2^ However, the actual process of identifying disease-causing variants and mutations remains a challenging task, and often one that requires at least some bioinformatics knowledge. This is due mainly to the sheer number of variants routinely identified in ES projects, the diversity of biological mechanisms by which variants may act, and the need to integrate large amounts of information from both pathogenicity scoring algorithms and clinical and population databases.

In this context, user-friendly graphical and interactive software tools have been developed that are able to filter, display and contextualise exome sequencing data in order to accelerate the discovery of disease-causing variants. These resources vary in the amount of public information they integrate, their interactivity, and the level of bioinformatics expertise required to execute them. For example, Genome Mining (GEMINI)^3^ allows the user to interactively explore their own variation files and overlaps information from dbSNP,^4^ ENCODE,^5^ ClinVar^6^ and KEGG,^7^ but requires users to have a good understanding of the command line and to be able to construct MySQL queries. Similarly, VCF-Miner^8^ and BrowseVCF^9^ allow the user to interactively filter their own variant call format (VCF) annotations through a web interface, but do not leverage information from other sources such as dbSNP and COSMIC.^10^ Other tools like BiERapp^11^ and exomeSuite ^12^ allow extensive variant filtering but do not support data visualisation. Similarly, tools that focus on variant visualisation at the protein level rather than filtering have also been developed, such as ProteinPaint,^13^ VizGVar^14^ and vcf.iobio.^15^ These resources are highly interactive, but either only display information already existing in databases without letting the user analyse their own variants (VizGVar: Ensembl),^16^ do not allow the user to see gene-level information (vcf.iobio) or require the user to perform several bioinformatics steps to see their data (ProteinPaint).

Here, we introduce VCF/Plotein, a user-friendly graphical web application to both visualise and filter variant information from exome sequencing studies that requires no bioinformatics skills or knowledge. As such, this application can be used by patients to explore their own genetic information, by biologists whose projects involve exome sequencing or by medical professionals studying a particular disease. VCF/Plotein allows the user to easily load a variant call format (VCF) file, identify genes with variants, and filter and visualise the variants in any gene with information about transcripts, protein domains, variant consequences and allelic frequencies in external databases. Furthermore, this application is especially suited for the medical community mindful of patient privacy, as it allows for sharing of selected data among collaborators while avoiding having to upload sample information to the server, instead running all operations locally. The resulting protein-level graphs are fully customisable, allowing the user to effortlessly generate professional vector images from their datasets for sharing, for presentations or for publication.

## METHODS

VCF/Plotein has been implemented entirely as a single-page application hosted on the Surge content-delivery network (CDN), to optimise static content downloading times across the globe. The application has been written mainly in JavaScript and uses the Vue.js-based Nuxt.js framework to control the storage, flow and presentation of information in the browser. A purpose-made API has been developed to obtain information from locally-installed external databases (gnomAD,^17^ dbSNP, COSMIC, ClinVar and GO term information for each annotated gene), and is running on a server with a 2-core Intel Xeon E5-4627 v4 2.60Ghz processor running a VMware 6.5.0 virtual machine over a Linux Centos 7.5 operating system. The server also has 4GB of RAM and a solid-state hard disk drive with 1TB of storage space. VCF/Plotein works with files in the VCF format, the standard format for genetic variant data storing, which consists of fields with the genomic position, reference and alternative base changes, ID, quality score, quality filters, other metrics including custom annotations and sample genotypes.^18^ Upon loading, a VCF is validated and its chromosome, position, reference and alternative fields are parsed, or an error is returned to the user. After identifying the assembly version from the appropriate line in the VCF, genes with variants are quickly found by matching an interval tree algorithm to the internal coordinate indexes containing each gene’s genomic positions. This generates a list with all the genes represented in the VCF, which can be filtered in different ways, such as by chromosome or by gene ontology [GO] term, in order to facilitate gene prioritisation. Once a gene is selected, information about protein-coding transcripts and functional domains is extracted from Ensembl via their REST API, independently of the existing custom annotations but without altering the original file. Dimensioning to a single gene is a key aspect for the performance of this application, since it considerably reduces computational load and information transfer over the internet. Consequences from all variants falling within the selected gene, as well as their pathogenicity scores by SIFT^19^ and PolyPhen,^20,21^ are obtained via the Ensembl Variant Effect Predictor.^22^ Cross-referencing with supported external databases is then performed via querying our internal database using the Elasticsearch search engine, which allows us to perform a smaller number of queries in an efficient manner. A flowchart depicting the process flow and system architecture of VCF/Plotein is shown in **Figure 1**. All collected information is stored as an array of objects in JSON format, returned to the web browser and depicted over a customisable plot of the primary structure of the canonical transcript (**Figure 2**). The protein graph is made using the D3.js library since it allows to easily map the information to vector graphic elements in the web page. All operations, except for the search of naked genomic positions in supported external databases, is performed locally using the user’s CPU.

**Figure 1.**
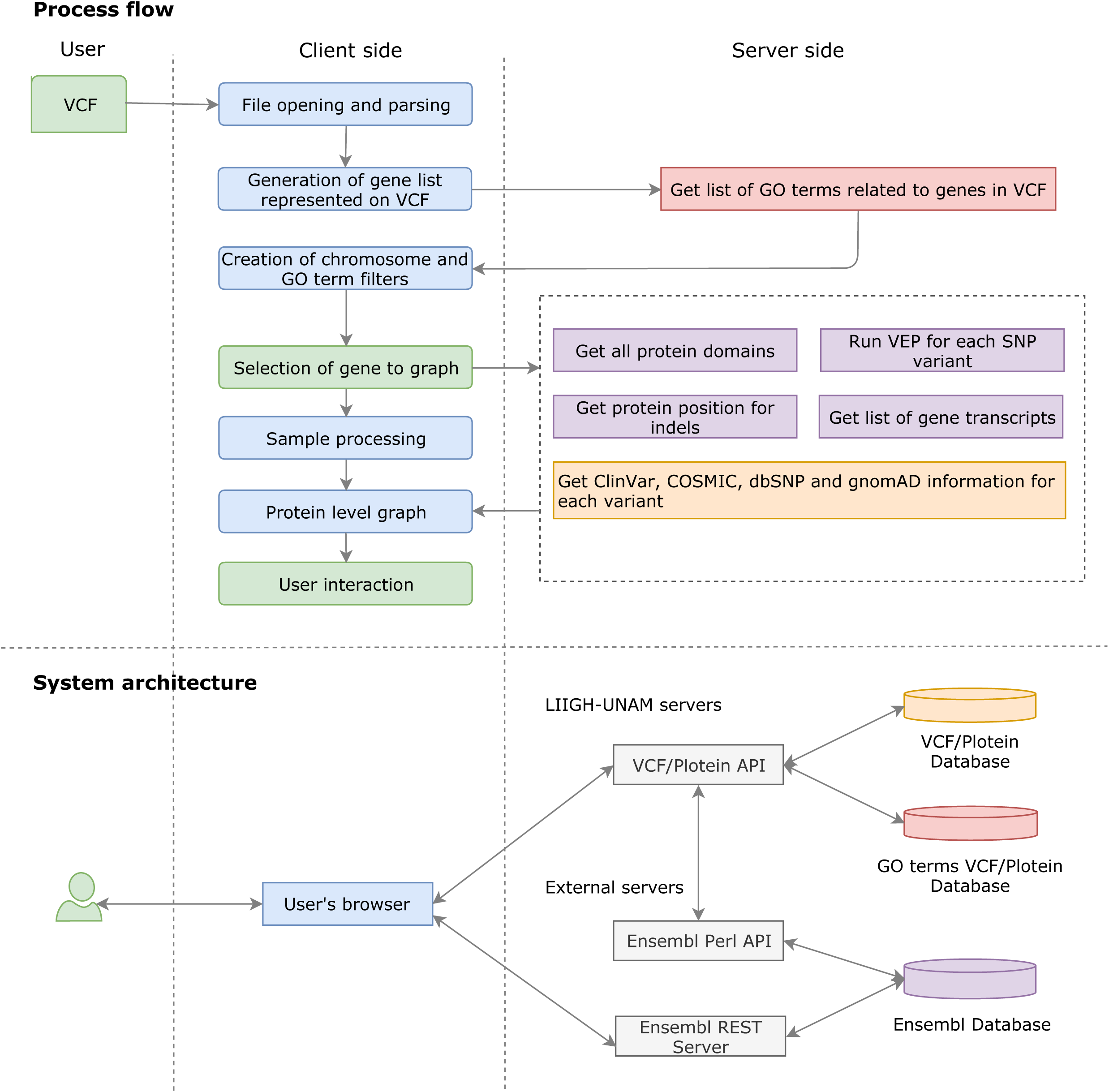
Chart depicting the VCF/Plotein process flow and system architecture. The colours of each box in the top panel correspond to the colour of the system architecture element in which it is executed (in the bottom panel), *e.g*., actions in green boxes are executed by the user, those in blue boxes are executed by the user’s browser, and actions in red, purple and orange are processes used to obtain information from diverse databases.

**Figure 2.**
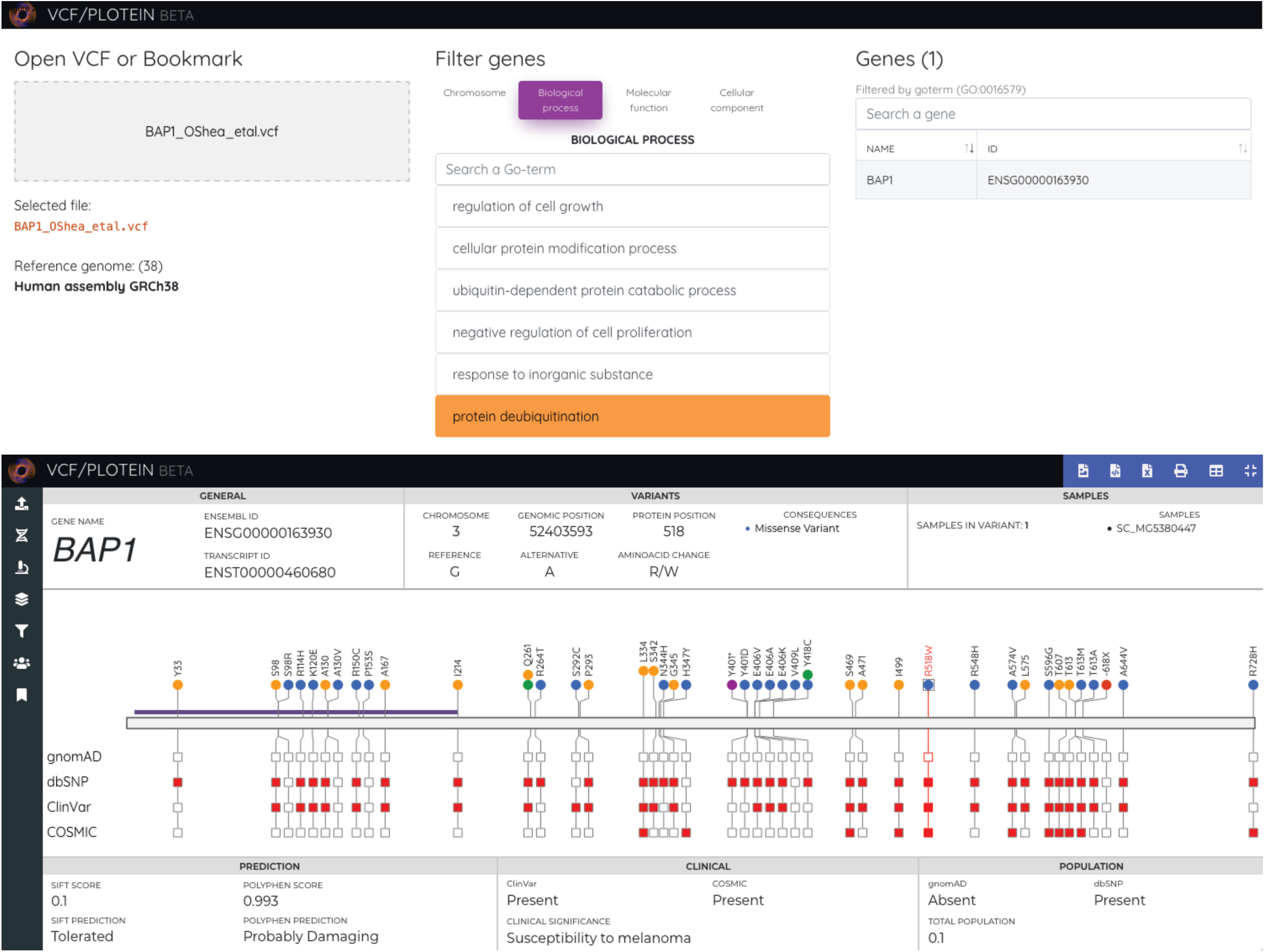
VCF/Plotein working screens. Top panel: Loading and data filtering screen. In the left column, the loaded VCF or bookmark is shown along the identified genome assembly. In the middle column, different gene filtering criteria are shown. The right column allows the user to select an individual gene to examine in detail. Bottom panel: Screen depicting protein-level information for the selected gene. The primary protein structure for the canonical transcript is shown alongside protein domains and other features, and the genetic variants falling on it are shown as lollipops. The colours of these indicate the protein consequence, and the squares below indicate the presence or absence of the variant in different external databases. Upon selection of a particular variant, other relevant data are shown in information bars at the top and bottom. Left-hand side and top menus allow the user to customise and interact with the plot. The depicted VCF contains variants published elsewhere (Ref. ^29^).

A menu, shown at the left-hand side of the browser, has been implemented to allow data filtering and graph customisation. The different features in this menu (Filters for transcript, functional domains and other protein features, variant type, variant frequency in external databases, variant pathogenicity score, variant clinical score and sample ID) all access information in the stored JSON object, which provides full interactivity due to the use of the Vue.js framework. A method has also been implemented that allows to export the current selected settings for any number of genes to a text file. This file can be used as a bookmark, which can be easily shared and loaded into VCF/Plotein for visualisation.

Functions to visualise and export the variant information in the customised plot to a CSV file, as well as to download and print the customised graph into a SVG or a PNG file, have also been implemented and are shown in a menu at the top right of the page.

## RESULTS

### Features

#### Overview

The only requirements to run VCF/Plotein are a computer with an internet connection and a VCF file. Once the user loads the VCF file, the genome assembly is identified, genes with variants are found, and a list of criteria is displayed to aid with gene prioritisation (by chromosome or by GO term category) (**Figure 2**, top panel). Once a gene is selected, a new page is shown with the primary protein structure of its canonical transcript with its domains and other features along with all its recorded variants. Variants are shown with an indication of their frequency among samples in the VCF, their transcript consequences, and their presence or absence in the gnomAD, dbSNP, ClinVar and COSMIC databases. The user can click on any variant to access further information about it, such as its genomic coordinates, a prediction of their pathogenicity according to SIFT and PolyPhen and a list of carrier samples (**Figure 2**, bottom panel). The left-hand menu allows the user to load a new VCF file, to select a different gene, to select a different transcript, to select which protein domains and features to show, to filter variants, to analyse sample IDs, and finally to bookmark the selected features. Using the top menu, variant information can also be displayed and downloaded in table format, as well as printed in the SVG vector image file format or the PNG raster graphics format.

#### Data security

The API and the internal databases have been installed behind a Fortinet firewall, and run over an HTTPS port with a SSL certificate for secure data transfer. No sensitive sample information is uploaded to the server, as queries to the internal databases consist only of naked genomic positions. Therefore, the server does not hold or save any sample information, an important feature given the data security policy that many patient-focused sequencing projects are bound by. All data processing, including construction of the JSON object and graphing of protein structures, is done locally.

#### Variant filtering and visualisation

Variants falling in any selected protein-coding transcript from any gene can be filtered and plotted. Users can filter variants by protein consequence (for example, to show only stop gains or splice site mutations), by clinical prediction (for example, to display only those present in the ClinVar or COSMIC datasets), by pathogenicity score (for example, only those scored as deleterious by SIFT and/or PolyPhen) or by their presence in allele frequency databases (for example, to display only variants not previously seen in gnomAD or dbSNP). Users can also select which protein domains and features to display (from InterProScan^23^ and Pfam^24^ and the ncoils,^25^ SEG,^26^ SignalP^27^ and TMHMM^28^ programs, as annotated by Ensembl), as well as customise the colours for each of these features. The customised protein plot can then be exported as an SVG or PNG file to use in presentations or
publications. Because of the nature of this web application, it is also well-suited to visualise information from large databases such as dbSNP or ClinVar.

#### Performance

VCF/Plotein is able to process VCF files from exome sequencing studies in a reduced time frame. We have tested our application with different file sizes in a number of system architectures and web browsers. The results are shown in **Table 1**.

**Table 1.** Performance of VCF/Plotein after loading three different VCF files in different operating systems with a range of hardware specifications. Tests were performed in the Google Chrome browser. All times are in milliseconds (ms).

|  | Computer specifications | Processes | Single gene VCF<br>Size: 2.9 mb<br>Number of genes: 1<br>Gene selected: 490 variants<br>Samples: 201 | ClinVar VCF<br>Size: 170.7 mb<br>Number of genes: 7357<br>Gene selected: 340 variants<br>Samples: NA | COSMIC VCF<br>Size: 421.8 mb<br>Number of genes: 23141<br>Gene selected: 1979 variants<br>Samples: NA |
| --- | --- | --- | --- | --- | --- |
| 1 | Operating system: macOS Mojave<br>Processor: 2.2 GHz intel core i7<br>RAM: 8 GB 1600MHz DDR3<br>Hard disk: 500 GB | File opening and parsing<br>Generation of list of genes<br>Creation of GO term filters<br>Fetching of variant information | 62<br>80<br>5<br>5724 | 1665<br>2821<br>1486<br>6499 | 4390<br>32300<br>11277<br>22192 |
| 2 | Operating system: Manjaro Linux<br>Processor: 1.7GHz AMD A8 x 4<br>RAM: 7 GiB<br>Hard disk: 750 GB | File opening and parsing<br>Generation of list of genes<br>Creation of GO term filters<br>Fetching of variant information | 222<br>16<br>255<br>6159 | 11528<br>4954<br>12566<br>31141 | 60492<br>75800<br>32418<br>106972 |
| 3 | Operating system: Windows 10<br>Processor: intel core i7-6500U<br>RAM: 8 GB 1600MHz DDR3<br>Hard disk: 1 TB | File opening and parsing<br>Generation of list of genes<br>Creation of GO term filters<br>Fetching of variant information | 18<br>32<br>5<br>4506 | 1033<br>1159<br>909<br>6754 | 10599<br>78363<br>10515<br>25364 |
| 4 | Operating system: macOS Mojave<br>Processor: 2.2 GHz intel core i7<br>RAM: 8 GB 1600MHz DDR3<br>Hard disk: 500 GB | File opening and parsing<br>Generation of list of genes<br>Creation of GO term filters<br>Fetching of variant information | 35<br>15<br>112<br>4974 | 2666<br>10648<br>10488<br>7042 | 33034<br>110945<br>29218<br>19251 |

#### Bookmarks

Bookmarks allow users to easily save any selected features from any number of genes in a text file (in JSON format) which can be easily shared so others can load it into VCF/Plotein and view and interact with this information. As only the information regarding the selected gene transcripts is saved, this function can be used by teams of clinicians and researchers studying a few candidate genes from exome-wide sequencing studies in order to facilitate data transfer and interpretation.

#### Comparison with other similar tools

VCF/Plotein combines variant filtering capabilities with an intuitive and interactive platform to visualise and customise protein plots. Furthermore, it does not upload any sensitive sample information to the server, instead running all operations locally. For these reasons, and because it requires no bioinformatics knowledge, we believe it is well-suited for patients exploring their own genetic data, for biologists analysing exome sequencing data, and for teams of scientists and medical professionals studying patients with a particular disease. Other available tools perform some of these functions, but either require at least some bioinformatics expertise, do not leverage information from external databases, or do not allow users to visualise their own exome data (**Table 2**).

**Table 2.**
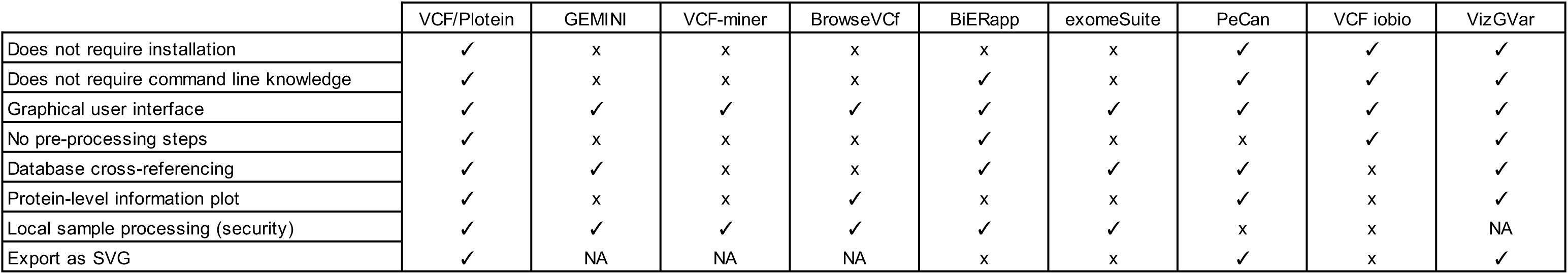
Comparison of the main features of VCF/Plotein with those of other similar tools.

|  | VCF/Plotein | GEMINI | VCF-miner | BrowseVCf | BiERapp | exomeSuite | PeCan | VCF iobio | VizGVar |
| --- | --- | --- | --- | --- | --- | --- | --- | --- | --- |
| Does not require installation | ✓ | x | x | x | x | x | ✓ | ✓ | ✓ |
| Does not require command line knowledge | ✓ | x | x | x | ✓ | x | ✓ | ✓ | ✓ |
| Graphical user interface | ✓ | ✓ | ✓ | ✓ | ✓ | ✓ | ✓ | ✓ | ✓ |
| No pre-processing steps | ✓ | x | x | x | ✓ | x | x | ✓ | ✓ |
| Database cross-referencing | ✓ | ✓ | x | x | ✓ | ✓ | ✓ | x | ✓ |
| Protein-level information plot | ✓ | x | x | ✓ | x | x | ✓ | x | ✓ |
| Local sample processing (security) | ✓ | ✓ | ✓ | ✓ | ✓ | ✓ | x | x | NA |
| Export as SVG | ✓ | NA | NA | NA | x | x | ✓ | x | ✓ |

## DISCUSSION

The ability to easily filter and display genomic information at the protein level is expected to contribute importantly to the identification of genetic variants conferring high-risk to develop various diseases by allowing non-bioinformaticians (e.g. many medical doctors and bench researchers) to perform variant prioritisation. Here, we have described VCF/Plotein, a novel web application capable of loading VCF files from exome sequencing projects in order to display, in a graphical and interactive manner, variants falling in any chosen transcript from any gene. Furthermore, variants can be filtered and customised to display only those of interest to the user, which can then be easily shared via a simple text file that can also be loaded into the web application. This kind of analyses can sometimes take an experimented bioinformatician hours or days (given the need to download databases, create custom scripts for filtering and find adequate tools for data visualisation), however here we have simplified this process and shortened the time required for analysis to minutes or seconds.

Our methodology also entails a number of advantages: By building VCF/Plotein as a single-page application, we are able to implement an iterative and incremental software development strategy, undertake constant improvements and regularly add new features. Furthermore, it allows us to easily keep our internal data structures up-to-date, importing the latest versions of external databases as they are released.

We anticipate that VCF/Plotein will allow researchers, especially in small labs, to focus on biology-relevant questions instead of having to learn to install software dependencies, learn to use variant-annotation and cross-referencing tools, and become familiar with the UNIX and/or MySQL command line. As direct-to-consumer genetic testing becomes even more accessible, we also believe that this application can be used by patients analysing their own genetic information. By combining variant filtering and annotation in a single graphical and interactive tool, we hope that variant prioritisation will become easier, faster and more intuitive.

## AVAILABILITY

VCF/Plotein is freely available at http://vcfplotein.liigh.unam.mx

## SUPPLEMENTARY DATA

No supplementary data

## ACKNOWLEDGEMENTS

This work received support from Alejandro de León and Carlos S. Flores of the Laboratorio Nacional de Visualización Científica Avanzada from the National Autonomous University of Mexico. We are thankful to Mamunur Rashid, Aravind Sankar, Carolina Castañeda Garcia and Patricia Basurto Lozada for testing the application, and to Xavier Soberón Mainero, Daniel Piñero Dalmau and Rafael Palacios for feedback during the development of this project. Finally, we would like to thank Abigayl Hernández and Eglee Lomelín for their continued support.

## FUNDING

This work was supported by the Wellcome Trust [204562/Z/16/Z to C.D.R.-E.] and Cancer Research UK [to D.J.A.] and Programa de Apoyo a Proyectos de Investigación e Innovación Tecnológica (PAPIIT UNAM) [IA200318 to C.D.R.-E, including an undergraduate scholarship to D.S.A.-M., and IA206817 to O.I.G.-S.]. R.O. is a PhD student from Programa de Doctorado en Ciencias Biomédicas, Universidad Nacional Autónoma de México (UNAM) and is supported by CONACyT [scholarship no. 573128]. Infrastructure for computational analyses was additionally provided by a CONACyT grant led by Dr. Alejandra Medina-Rivera [269449].

## Notes

**CONFLICT OF INTEREST:** No conflicts of interest.

